# Nascent adhesions differentially regulate lamellipodium velocity and persistence

**DOI:** 10.1101/2021.11.15.468602

**Authors:** Keith R. Carney, Akib M. Khan, Shiela C. Samson, Nikhil Mittal, Sangyoon J. Han, Michelle C. Mendoza, Tamara C. Bidone

## Abstract

Cell migration is essential to physiological and pathological biology. Migration is driven by the motion of a leading edge, in which actin polymerization pushes against the edge and adhesions transmit traction to the substrate while membrane tension increases. How the actin and adhesions synergistically control edge protrusion remains elusive. We addressed this question by developing a computational model in which the Brownian ratchet mechanism governs actin filament polymerization against the membrane and the molecular clutch mechanism governs adhesion to the substrate (BR-MC model). Our model predicted that actin polymerization is the most significant driver of protrusion, as actin had a greater effect on protrusion than adhesion assembly. Increasing the lifetime of nascent adhesions also enhanced velocity, but decreased the protrusion’s motional persistence, because filaments maintained against the cell edge ceased polymerizing as membrane tension increased. We confirmed the model predictions with measurement of adhesion lifetime and edge motion in migrating cells. Adhesions with longer lifetime were associated with faster protrusion velocity and shorter persistence. Experimentally increasing adhesion lifetime increased velocity but decreased persistence. We propose a mechanism for actin polymerization-driven, adhesion-dependent protrusion in which balanced nascent adhesion assembly and lifetime generates protrusions with the power and persistence to drive migration.

## Introduction

Cell migration emerges from the controlled assembly of macromolecules that generate the mechanical forces of cell migration. Directional migration is specifically associated with the velocity and persistence of outward motion of the leading edge, termed lamellipodium protrusion [1, 2]. The edge protrusion is generated by actin filaments polymerizing against the plasma membrane and the formation of adhesions to the substrate [3, 4]. Actin polymerization generates pushing force that overcomes membrane tension and moves the cell edge forward [3, 5–7]. Counterforce from the membrane simultaneously induces actin retrograde flow away from the membrane and towards the cell center [7]. The actin retrograde flow becomes physically anchored to the substrate by adhesions, which transmit the flow to the substrate as traction force [8, 9]. Adhesion-mediated traction force promotes edge protrusion, but induces edge retraction if excessive [8, 10]. The pushing and traction forces are balanced for protrusive activity, but how the individual molecular dynamics generate this balance remains unknown.

Actin polymerization against the plasma membrane has been described as a stochastic Brownian ratchet [11]. In the Brownian ratchet model, fluctuations in the membrane and the actin filaments that abut the membrane create gaps between the two structures, which allow for the addition of new monomers that push against the leading edge [11]. The lamellipodium harbors an excess of monomeric actin that polymerizes onto existing filaments [7]. The actin nucleator ARP2/3 increases net actin polymerization by initiating new filaments off of the sides of existing filaments [12, 13]. Thus, ARP2/3 increases the number of actin filaments abutting and ratcheting against the membrane.

Fibroblasts and most epithelial cells move via cycles of edge protrusion, in which protrusions progress through protrusion initiation, reinforcement, and retraction phases that result from changes in membrane tension [14, 15]. Protrusions are initiated by the un-tethering of actin filaments from the membrane [16–18]. As the protrusion progresses, the membrane is stretched and membrane tension increases, which pushes back on the actin filaments and decreases the likelihood of new monomer addition [14, 19–21]. The mechanism by which filament elongation decreases with tension is explained by a force-velocity relationship in which actin filament elongation stalls at high force [22]. ARP2/3 activity increases after protrusion initiation in the reinforcement phase, which is also named the power phase in reference to the increased number of actin filaments pushing against the membrane [23–26]. Despite the increased pushing force, the mechanical feedbacks between membrane tension and actin polymerization cause edge velocity to slow after protrusion initiation.

The formation of small, transient adhesions in the lamellipodium, termed nascent adhesions, promotes and is required for protrusion velocity and persistence in protrusion-retraction cycles [27, 28]. Nascent adhesions work a molecular clutch, engaging and anchoring the actin filaments undergoing retrograde flow to the extracellular matrix on the substrate [3]. This decreases actin flow velocity [14, 29] and converts the motion into traction on the substrate and cell edge motion [30]. Because anchoring actin filaments onto adhesions also maintains the actin barbed ends against the membrane, the assembly rate of nascent adhesions correlates with leading edge velocity [31]. Molecular clutch disassembly depends on the amount of traction on the adhesions [14, 32]. Adhesion traction peaks along with the rate of actin retrograde flow in the reinforcement phase of edge protrusion [14, 24, 33]. The lifetime of nascent adhesions initially increases along with actin flow, but then decreases as actin flow increases further [10, 24], resulting in a stick and slip mechanism between actin filaments and adhesions. When nascent adhesions disassemble, they disengage from both the substrate and actin filaments [10].

The difficulty in experimentally isolating actin polymerization without affecting adhesion dynamics and vice versa has precluded a complete understanding of how lamellipodial actin polymerization and adhesion assembly and disassembly work together to control lamellipodium protrusion. In order to assess the contributions of actin and adhesion dynamics to edge motion, we developed a computational model that incorporates the Brownian ratchet mechanism (BR) and the molecular clutch mechanism (MC) [10, 34]. The model revealed that increasing the lifetime of adhesions supports cell edge velocity, but it reduces motional persistence in the initial phase of protrusion. Manipulation and computerized tracking of adhesion lifetime in live cells confirmed that adhesion lifetime promotes protrusion velocity but decreases motional persistence. Together, our findings suggest a previously unappreciated role for nascent adhesion force-dependent clutch mechanism in the control of initial leading-edge motion.

## Results

We developed a novel computational model of lamellipodium protrusion based on Brownian dynamics. The model incorporates: explicit actin filaments represented as polar rods of interconnected units; nascent adhesions represented as dynamic point particles that link filaments to a fixed substrate; and a flexible membrane represented as a series of rigid rods connected by springs (Figure 1). We designed a 2D domain of 2 μm x 0.5 μm size, in which actin filaments fluctuate under thermal motion, polymerize, branch and depolymerize, and link to adhesions. Actin polymerization produces a force against the membrane (***F***_*Pol*_), which pushes the membrane forward. The displacement of the membrane produces an increase in membrane tension (***F***_*M*_), which pushes the filaments away from the membrane and results in actin retrograde flow. ***F***_*M*_ also decreases the actin polymerization rate according to the decreasing exponential force-velocity relation characteristic of the Brownian ratchet (BR) mechanism [11, 22, 35–37]. Filaments can link to substrate adhesions, which convert their retrograde flow into adhesion traction (***F***_*A*_). The model ensures force balance between ***F***_*Pol*_, ***F***_*M*_, and ***F***_*A*_, resulting in relative movements of filaments and membrane at every time step (Figure 1A). The model simultaneously implements the Brownian ratchet mechanism of actin filament polymerization (BR) and the molecular clutch mechanism of adhesion dynamics (MC). The BR is implemented through the force-dependent velocity of filament elongation under membrane tension [11, 22, 35–37] (Figure 1B). The MC is implemented with biphasic force-dependent unbinding of adhesion clutches [38, 39] (Figure 1C).

**Figure 1.**
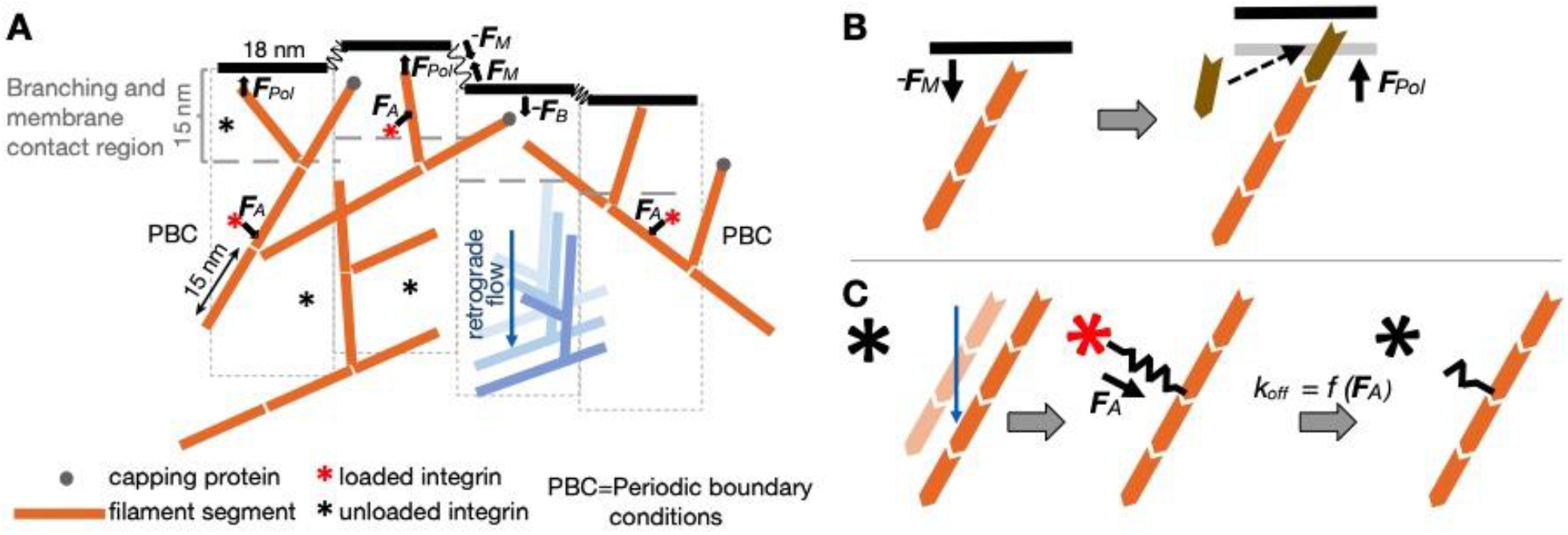
BR-MC model of lamellipodium protrusion. (**A)** Elements and interactions in the 2D computational model. Actin filaments are represented as polarized rigid rods of consecutive 15 nm units with each unit representing 5 actin monomers. Capping proteins bind the filaments with rate *k_cap_* and prevent further polymerization. Each actin filament can form branches at a 70° angle within 15 nm from the edge membrane (*k*_*branch*_). Polymerization with rate *k_pol_* against the membrane pushes the membrane forward (***F****_Pol_)*. The membrane is represented as rods interconnected by harmonic potential energies with defined stiffness *k*_*m*_, which create tension upon displacement, ***F***_*M*_. A boundary force (***F***_*B*_) prevents the filaments from crossing the membrane. The membrane and boundary forces restrain actin polymerization and induce retrograde flow. Adhesions are represented as integrins that undergo cycles of activation and deactivation, with *k*_*on*_ and *k_off_,* respectively. When active (black star), the adhesions connect actin filaments if within a proximity of 15 nm (red star). Adhesion engagement is represented as a spring and builds tension against the substrate in response to actin retrograde flow (adhesion tension, or traction force, ***F***_*A*_). Adhesion density is maintained within the physiological range ~500 integrins/μm^2^ [77]. A periodic boundary condition wraps the model edges and creates an infinite domain in which all the membrane segments experience spring tension on both sides. For each time step, the new position for each actin filament is computed from the sum of Brownian forces, ***F***_*B*_, ***F***_*M*_, and ***F***_*A*_. The new position for each membrane segment is computed from the sum of ***F***_*Pol*_, ***F***_*M*_, and ***F***_*A*_. (**B)** The BR mechanism. Polymerized actin filaments (orange) undergo thermal motion in the 2D domain near the membrane. Addition of a new monomer (brown) against the membrane pushes the membrane outward with ***F***_*Pol*_. (**C**) The MC mechanism. Actin undergoes retrograde flow (blue arrow) and integrins are activated (black star). Activated integrins bind to an actin filament to create adhesions with traction force (***F***_*A*_, red star) that slows the filament’s flow and controls adhesion unbinding.

### The BR-MC model reproduces physiological lamellipodium motion

The BR-MC model produces continuous filament polymerization, filament branching against the membrane, and nascent adhesion assembly and disassembly. Membrane displacement, increased membrane tension after displacement, actin retrograde flow, and adhesion-mediated anchoring of the flow emerge from the molecular dynamics (Figure 2A and Figure 2 - movie supplement 1). We first tested that our model reproduces previous experimentally measured properties of the lamellipodium: edge motion velocity, actin retrograde flow velocity, and adhesion traction force [33, 40]. In order to verify how actin filament polymerization alone affects edge motion, we initially set membrane tension to *k*_*m*_ = 0.3 pN/nm, which is a value consistent with measurements in epithelial cells [41, 42]. We used a constant rate of actin filament branching (*k*_*branch*_ = 0.5 s^−1^) that corresponds to experimentally measured ARP2/3 activation [43], and a constant rate of adhesion unbinding (*k*_*off*_ = 0.1 s^−1^) that matches the unloaded lifetime of integrin-ligand bonds [44–46]. We found that individual membrane segments exhibited initial velocity peaks of 30 - 40 nm/s and slower velocities as the protrusion progressed (Figure 2B). Within the corresponding segments of the lamellipodium, actin retrograde flow increased as the protrusion progressed (Figure 2C). Traction forces exhibited peaks of 60 - 80 pN (Figure 2D), consistent with experimental measurements of individual adhesions [32]. Plotting membrane and actin retrograde flow velocities as a function of traction force showed that membrane velocity increased and actin retrograde flow decreased as traction increased from 0 to 40 pN (Figure 2E), as previously described [30, 32].

**Figure 2.**
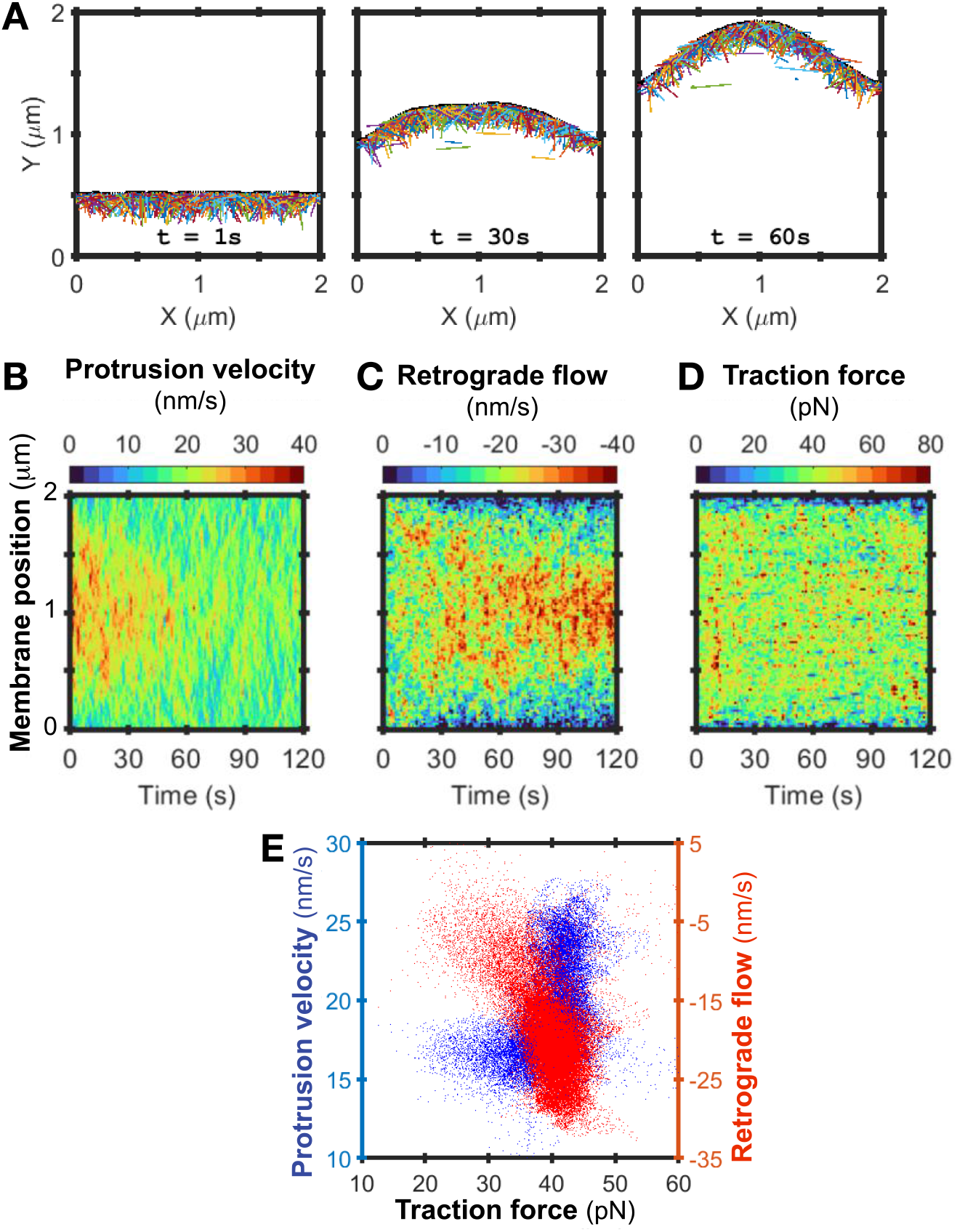
Model reproduces physiological rates of edge protrusion and retrograde flow. (**A**) Snapshots from model simulations over run times in seconds (s). Individual filaments are depicted in different colors. Parameters listed in Table 1. Membrane movement in Y from the sum of forces (*F*_*pol*_, *F*_*M*_, and *F*_*A*_). (**B**) Heatmap of mean edge velocity at each 20 nm membrane position (18 nm rod + 1 nm of spring on each side). 2 minute (min) simulation with *k*_*m*_ = 0.3 pN/nm, *k*_*branch*_ = 0.5 s^−1^, *k*_*on*_ = 1 s^−1^, *k*_*off*_ = 0.1 s^−1^, and periodic boundary conditions. Sampling every 0.001 s, averaged over 1 s intervals. Representative of *n*=10 iterations. (**C**) Heatmaps of retrograde flow and (**D**) traction force, computed from simulations, sampling, and averaging as in B, and smoothening using a 2-D Gaussian filter with SD = 0.5. (**E**) Edge protrusion velocity and actin retrograde flow plotted as a function of traction force. Data from 3 model runs under conditions in B, but smoothened using a Gaussian smoothing kernel with standard deviation (SD) = 2. Points are values for individual membrane segments. Retrograde flow values within +/- 1000 nm/s were plotted, ≥ 94% of values.

We averaged the velocities and traction forces across the middle 1 *μm* membrane segment to mimic the computerized measurements from experimental images limited by the resolution of light microscopy. The mean protrusion velocity peaked at 24.3 nm/s at 10 s and slowed to 17.0 nm/s at 120 s (Figure 2 - figure supplement 1A), consistent with published measurements of PtK1 epithelial cells using (~25 nm/s [24, 33]). Actin retrograde flow increased after protrusion initiation, from a mean of 18.9 nm/s at 10 s to a mean of 26.7 nm/s at 120 s (Figure 2C and Figure 2 - figure supplement 1B), also consistent with PtK1 measurements (11.7 - 21.7 nm/s [24, 33]). The differences in time at which maximum edge velocity and maximum retrograde flow were reached are in quantitative agreement with the observed timing of lamellipodia actin retrograde in areas lacking significant traction force [24, 32, 33, 40].

**Table 1.**
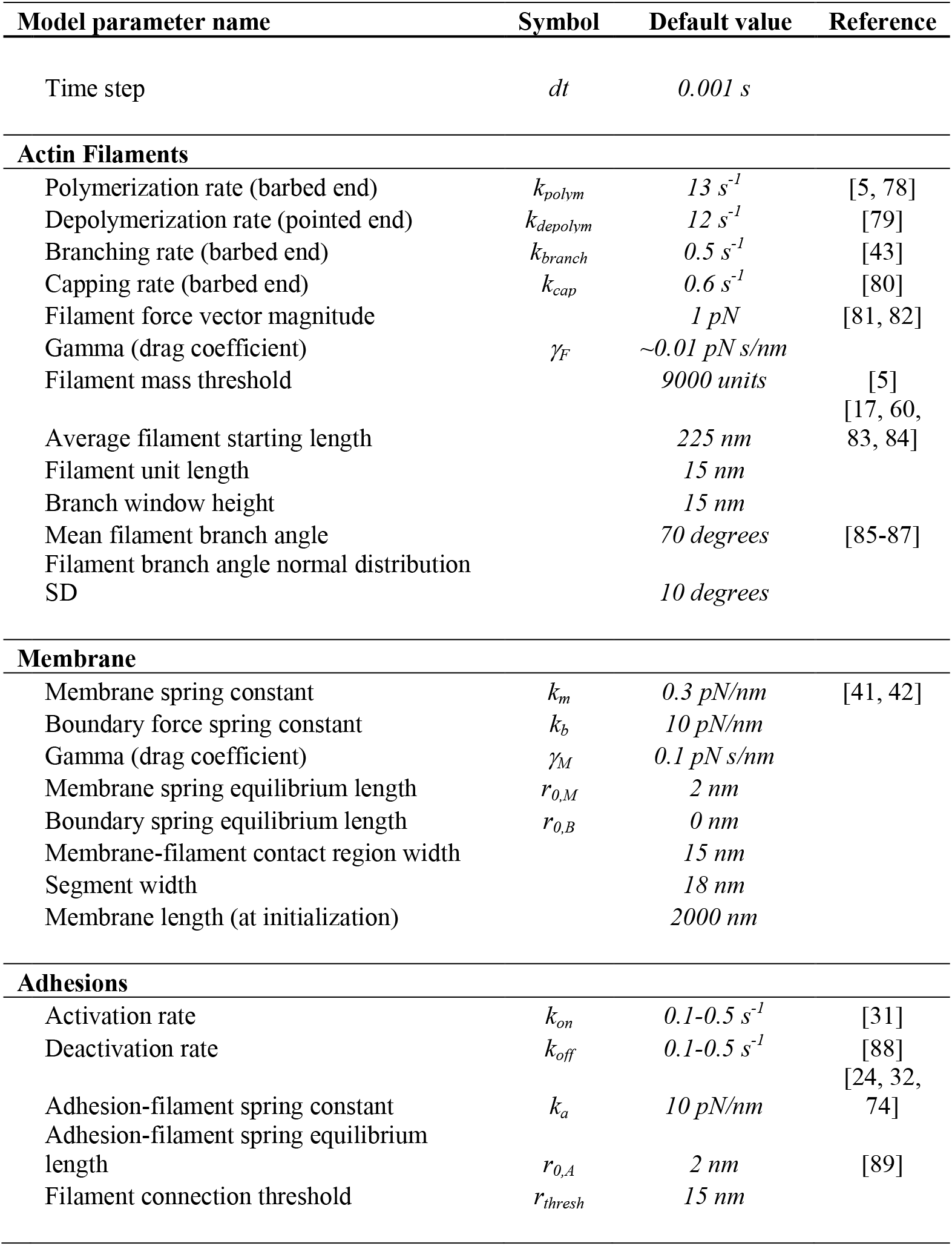
List of model parameters.

We also validated the effects of membrane tension on the force relationships. In order to capture a greater range of edge velocity values, we modeled the nonmotile portion of membrane in cells by fixing the side boundaries of the modeling domain. Similar to our observations with unfixed edges, edge protrusion slowed and retrograde flow increased as the protrusion progressed (Figure 2 –figure supplement 2A-B). Traction forces were stable, but high at the edges due to the high concentration of actin filaments and adhesions in the unstretched, tethered space (Figure 2 –figure supplement 2C). We ran the model with low membrane tension (0.03 pN/nm) to replicate the scenario of protrusion initiation and with high membrane tension (3 pN/nm) to represent the protrusion reinforcement phase [15, 30, 47]). We found that under low membrane tension, actin retrograde flow positively correlated with protrusion velocity (Figure 2 –figure supplement 2D). In contrast, under high membrane tension, actin retrograde flow negatively correlated with protrusion velocity (Figure 2 – figure supplement 2D). This is consistent with experimental data that shows high retrograde flow from the high rate of actin polymerization against the tense membrane, but slow protrusion velocity due to the high tension [20, 31, 32, 52].

### Actin polymerization and adhesion assembly and disassembly control lamellipodium velocity

In order to decipher the contributions of actin filament polymerization and adhesion dynamics to membrane velocity, we systematically varied the rates of actin branching, adhesion assembly, and adhesion disassembly (*k*_*branch*_, *k*_*on*_, and *k*_*off*_). For simplicity, we averaged the velocity of the 101 membrane segments, including the fixed edges of the modeling domain, which resulted in lower velocities than reported in Figure 2. We found that increasing *k*_*branch*_ four-fold (from 0.2 to 0.8 s^−1^), with fixed adhesion assembly and disassembly rates (*k*_*on*_ = *k*_*off*_, = 0.1 s^−1^), resulted in a 30% increase in protrusion velocity (from 16.6 nm/s to 21.5 nm/s, Figure 3A). Increasing adhesion assembly rate *k*_*on*_ five-fold (from 0.1 to 0.5 s^−1^), with fixed *k*_*branch*_ = 0.5 s^−1^ and adhesion *k*_*off*_ = 0.3 s^−1^, resulted in a 10% increase in protrusion velocity (from 17.8 nm/s to 19.6 nm/s, Figure 3B). Increasing the disassembly rate of adhesions five-fold decreased protrusion velocity 10% (from 19.8 nm/s to 18.4 nm/s, Figure 3C). Together, these results indicate that actin assembly is the main driver of edge velocity and that actin’s control can be augmented by adhesion formation and lifetime. We tested how actin filament polymerization and adhesion dynamics together govern membrane motion by systematically varying *k*_*branch*_ and *k*_*off*_. Membrane velocity increased with *k*_*branch*_ and decreased with *k*_*off*_ (Figure 3D). We also tested how adhesion formation and lifetime control protrusion velocity by simultaneously varying *k*_*on*_ and *k*_*off*_ and found that protrusion velocity peaks with the highest adhesion assembly rate and lowest adhesion disassembly rate (Figure 3E). This indicates that protrusion velocity depends on both adhesion assembly and maintenance. We found that the traction force transmitted by adhesions also peaked with the highest adhesion assembly rate and lowest adhesion disassembly rate, suggesting that adhesion traction force promotes edge protrusion (Figure 3F).

**Figure 3.**
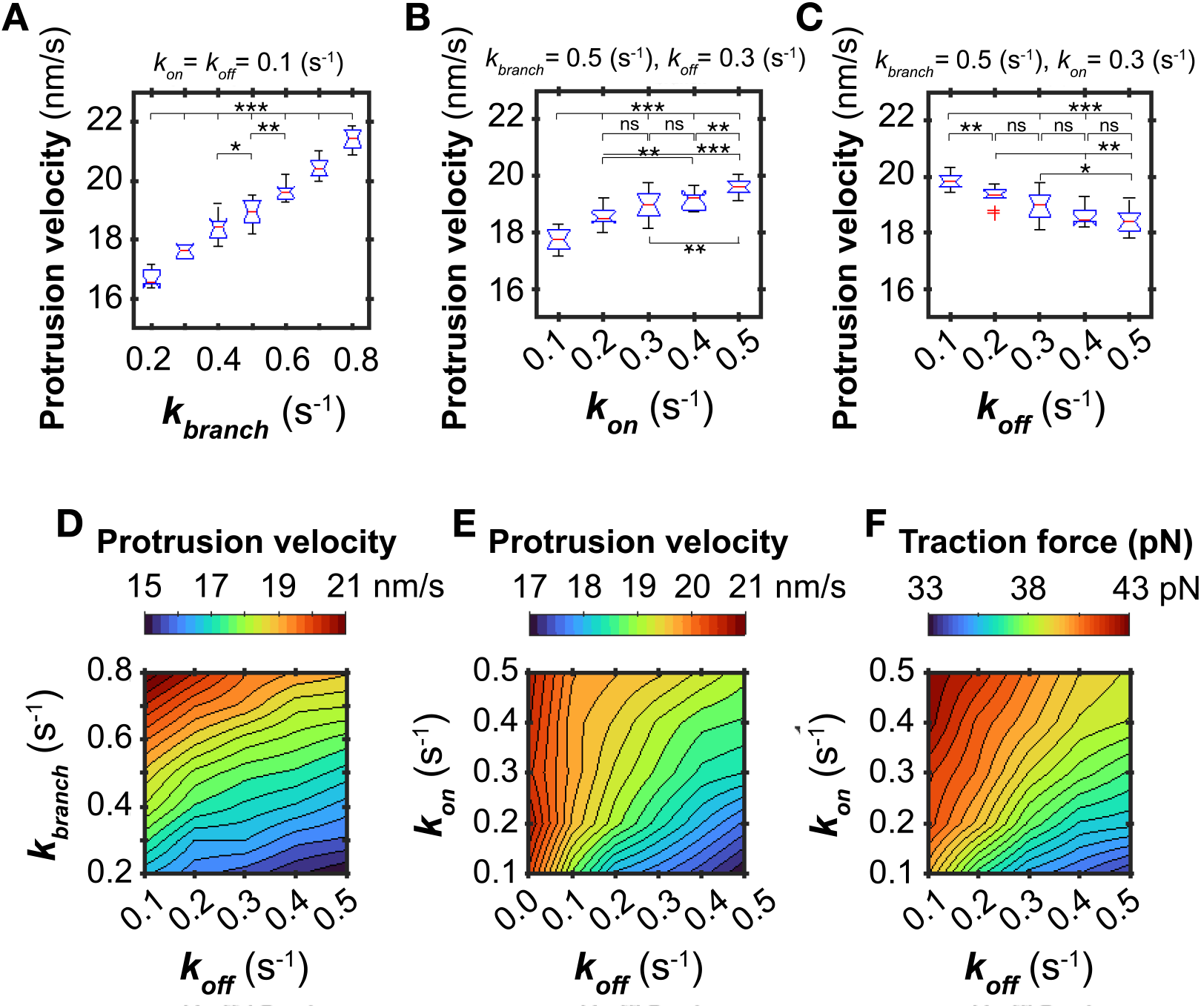
Actin assembly more significantly promotes protrusion velocity than adhesion assembly. (**A-C**) Distribution of lamellipodium velocity as a function of actin assembly rate *k*_*branch*_, adhesion *k*_*on*_ and adhesion *k*_*off*_. 10 s simulations with *k*_*m*_ = 0.3 pN/nm, maximum allowed adhesions = 1000 and periodic boundaries. Box plots’ red line is median velocity and notches 95% CI, from *n* = 10 simulations. Significance is Mann-Whitney U test of mean. Adhesion density ranged around 500, (A) 493 - 505/μm^2^, (B) 246 - 626/μm^2^, (C) 758 - 378/μm^2^. (**D**) Heatmap of mean protrusion velocity that results from variable actin *k*_*branch*_, adhesion *k*_*on*_ = 0.1 s^-^1, and variable adhesion *k*_*off*_. (**E**) Mean protrusion velocity resulting from *k*_*branch*_ = 0.5 s^−1^ and variable *k*_*on*_ and *k*_*off*_. (**F**) Mean traction force resulting from *k*_*branch*_ = 0.5 s^−1^ and variable *k*_*on*_ and *k*_*off*_. Force computed from identical conditions as in A-E, except simulations were 20 s.

### The lifetime of nascent adhesions regulates the velocity of the membrane

Adhesion traction determines the lifetime (τ) of adhesions, which is the inverse of unbinding rate (τ = 1/*k*_*off*_). To test the role of adhesion lifetime in protrusion, we incorporated a force-dependent molecular clutch mechanism into the computational model. The molecular clutch bases the probability of adhesion-actin bond breakage on tension and thus creates a variable rate of adhesion inactivation (*k*_*off*_). We tested three different force-lifetime relations for the adhesions, varying for maximum lifetime, τ_*max*_ (Figure 4A). The breaking point for the actin-adhesion bond, or force corresponding to τ_*max*_, was set at 30 pN [48–50]. The peaks in lifetime were either 3 s, 7.5 s, or 12 s, typical behavior of integrin unbinding from fibronectin under load [48]. We used these force-lifetime relationships to control the probability of adhesion-actin bond breakage during simulations with variable actin polymerization rate. We found that membrane velocity increased proportionally with both *k*_*branch*_ and τ_*max*_ (Figure 4B). Increasing actin assembly 4-fold resulted in a 17% increase in edge velocity (from 13.3 to 15.6 nm/s, using τ_*max*_ = 3 s), while increasing adhesion lifetime 4-fold resulted in about a 6% increase in velocity (from 15.6 to 16.5 nm/s, using *k*_*branch*_= 0.8 s^−1^). This result matched our findings on actin and adhesion-mediated control of edge motion in the absence of the molecular clutch (Figure 3D).

**Figure 4.**
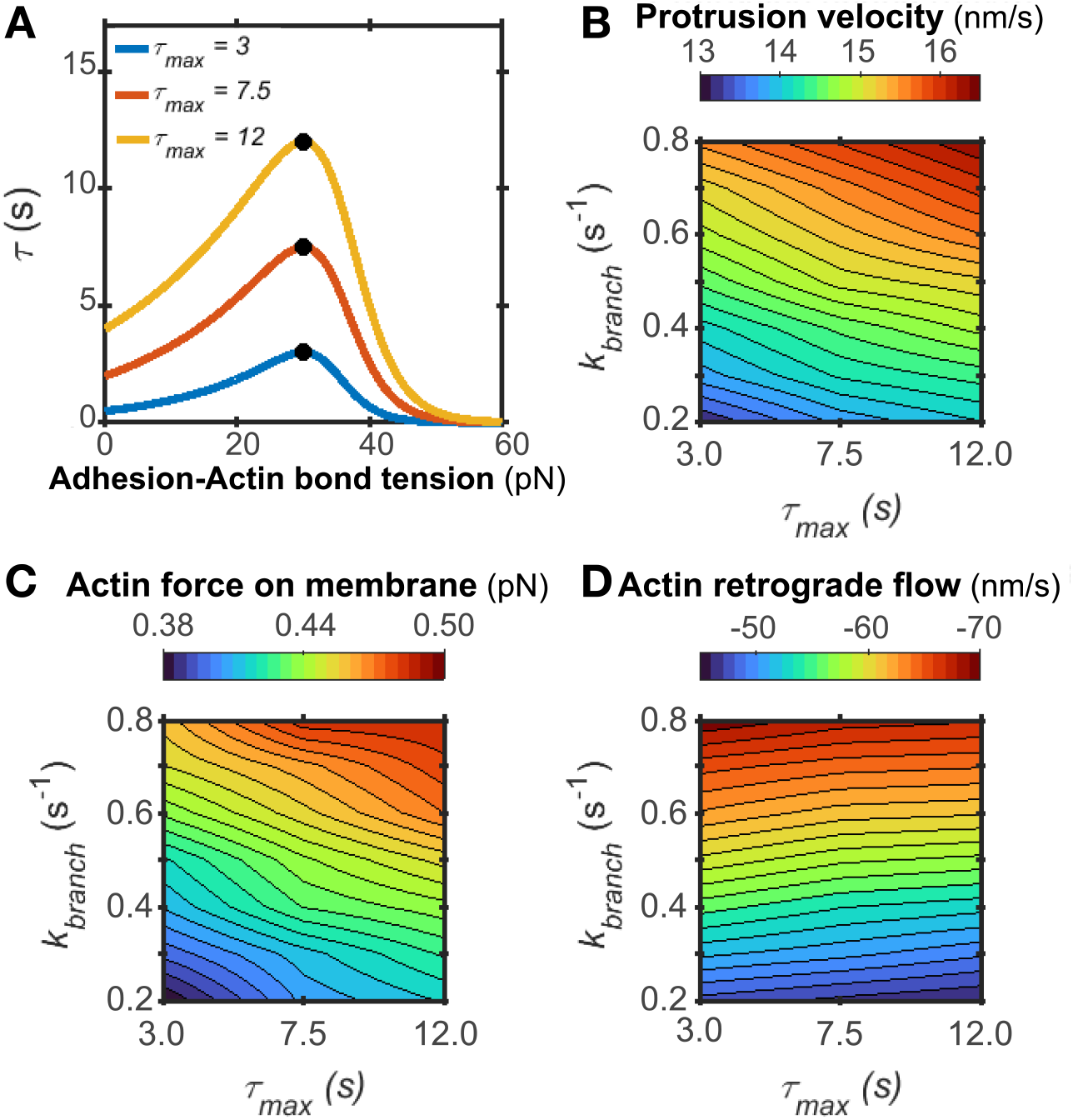
The molecular clutch mechanism contributes to leading edge motion. (**A**) Force-dependent relationship of the molecular clutch: adhesion lifetime (*τ*) versus adhesion-filament bond tension where *τ* = *1/k*_*off*_. Under high tension (50 pN), *k*_*off*_ = 2 s^−1^ and *k*_*off*_ increases exponentially with increased tension. (**B-D**) Mean protrusion velocity, actin force on membrane, and actin retrograde flow velocity that results from variations in *τ*_*max*_ and *k*_*branch*_. 10 s simulations with *k*_*m*_ = 0.3 pN/nm*, k_on_* = 1.0 s^−1^, molecular clutch engaged with *τ*_*max*_ = 3.0, 7.5, and 12.0 s with intermediate values interpolated, maximum allowed adhesions = 2000, boundaries fixed, *n* = 10. Because inactivated adhesions are removed from the modeling domain, *k*_*on*_ and the number of allowed adhesions were increased to obtain 500 adhesions/μm^2^ [77]. Membrane force calculated as the mean membrane spring tension for all membrane springs and all time points. All flow velocities are plotted.

Increasing adhesion lifetime can have the secondary effect of increasing adhesion density, observed in Figure 3. Using *k*_*branch*_ = 0.5 s^−1^, we found that increasing adhesion lifetime from 3 s to 12 s increased mean adhesion density from 299 to 505/μm^2^ (Figure 4 – Figure Supplement 1A). We identified the range of allowed adhesions in simulations with τ_*max*_ = 3 s that would generate adhesion density of ~300 to 500/μm^2^ (Figure 4 – Figure Supplement 1B). We then varied the maximum allowed adhesions to test how increasing adhesion density alone affects protrusion velocity, using the intermediate *k*_*branch*_ = 0.5 s^−1^. We found that increasing mean adhesion density from 310 to 513/μm^2^, independent of adhesion lifetime, increased protrusion velocity 2.3%, from 14.51 nm/s to 14.85 nm/s (Figure 4 – Figure Supplement 1C and D). Under the same conditions, increasing adhesion lifetime from 3 to 12 s resulted in a 4.6% increase in protrusion velocity, from 14.55 nm/s to 15.22 nm/s (Figure 4B and Figure 4 – Figure Supplement 1D). Thus, increased adhesion density resulting from increased lifetime does not fully explain adhesion lifetime’s effects on edge velocity. These findings substantiate our conclusions that actin polymerization rate is the most significant driver of edge velocity and adhesion lifetime augments the velocity.

We evaluated the contribution of actin assembly and adhesion lifetime to pushing force on the membrane and the retrograde flow that results from membrane counterforce. We found that force on the membrane increased with both *k*_*branch*_ and τ_*max*_, as peak force on the membrane occurred at the highest *k*_*branch*_ and τ_*max*_ (Figure 4C). Actin assembly was the main driver. With τ_*max*_ = 3 s, a four-fold increase in *k*_*branch*_ (from 0.2 – 0.4 s^−1^) increased pushing force 21% (from 0.38 to 0.46 pN). With *k*_*branch*_ = 0.2 s^−1^, four-fold increase in τ_*max*_ (from 3 – 12 s) increased pushing force 11% (from 0.38 to 0.42 pN). In contrast, actin retrograde flow increased with *k*_*branch*_ but decreased with τ_*max*_ (Figure 4D). The reduction in retrograde flow with adhesion lifetime is expected from previous experimental observations [14, 29]. These simulations with the τ_*max*_ variable suggest a critical role for the lifetime of nascent adhesions in regulating force against the membrane and the resulting edge velocity and actin retrograde flow.

### The lifetime of nascent adhesions controls the persistence of membrane motion

Because pro-migratory signaling pathways promote the disassembly of nascent adhesions [53], which limited protrusion velocity in our model, we hypothesized that adhesion disassembly might promote the alternative edge protrusion output of protrusion persistence. We quantified protrusion persistence as the time between the large oscillations in membrane velocity, determined by Empirical Mode Decomposition (Figure 5 – figure supplement 1). While adhesion lifetime promoted protrusion velocity at low membrane tension (0.03 pN and 0.3 pN, Figure 5A and B), it decreased its motional persistence (Figure 5C and D). Under high tension, adhesion lifetime promoted membrane velocity without inhibiting persistence (Figure 5E and F). We tested the hypothesis that increasing the lifetime of nascent adhesions reduces protrusion persistence by slowing the motion of actin filaments and shortening the time it takes for actin polymerization to stall. We computed the motion of actin filaments within 1 micron of adhesions and varying τ_*max*_. Increasing τ_*max*_ decreased the actin filaments’ mean displacements (Figure 5 – figure supplement 2A). Similarly, the time to stall, or decay time, for actin polymerization decreased with increasing τ_*max*_ (Figure 5 – figure supplement 2B). Thus, adhesion lifetime can decrease protrusion persistence by controlling actin filament mobility and polymerization.

**Figure 5.**
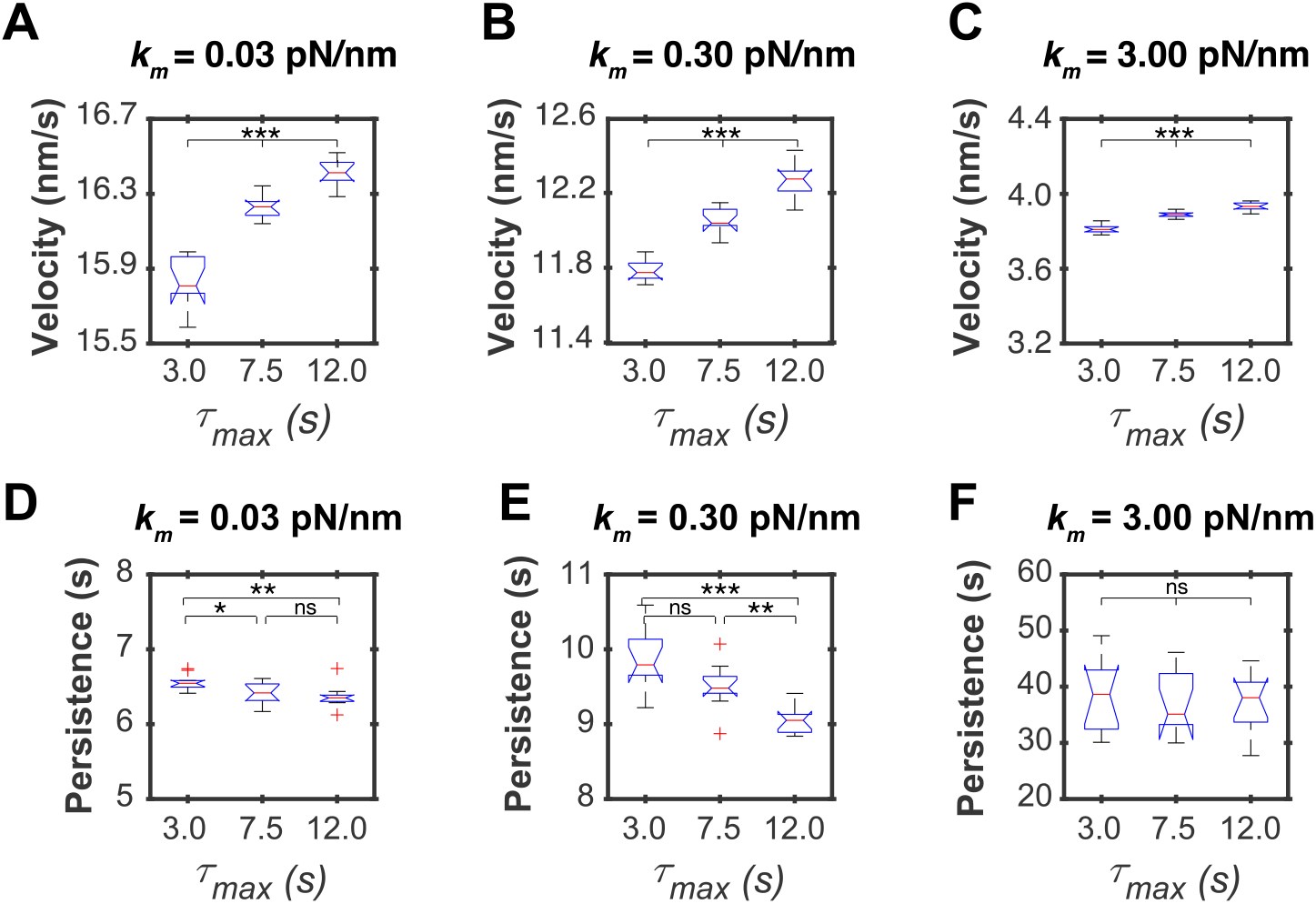
Adhesion lifetime promotes lamellipodia velocity but inhibits persistence. Model simulations in which membrane tension and molecular clutch *τ*_*max*_ varied, *k*_*branch*_ = 0.5 s^−1^, *k*_*on*_ = 1.0 s^−1^, boundaries fixed, 2 min run time. (**A-C**) Mean velocity measurements of the middle micron region of the membrane for each *τ*_*max*_ and each membrane tension. (**D-F**) Persistence measurements of the middle micron region of the membrane versus *τ*_*max*_ for each membrane tension. Persistence defined as the time between minimums in membrane segment velocities with minimums ≥ 5 nm/s. Box plots with red line marking medians and notches 95% CI. Significance is Mann-Whitney U test of means from *n* = 10 simulations.

### Integrin activation promotes lamellipodium velocity and decreases its persistence

To experimentally test the finding that adhesion lifetime promotes lamellipodia protrusion velocity but limits persistence, we labeled adhesions in COS7 epithelial cells using transient expression of Paxillin-mApple and imaged adhesion and edge dynamics during 5 min of steady-state migration. We segmented the adhesions using focal adhesion analysis software for quantification of the adhesions’ lifetime [51] and used morphodynamics software to track the edge motion [52] (Figure 6A and B). We noted that protrusions exhibited adhesions with heterogeneous lifetimes, in which clusters of short-living adhesions co-resided with a few longer-lifetime adhesions. The range of long lifetimes varied per movie, which appeared to be related with edge protrusion. For example, a cell in which the longest lifetimes are ~4.7 min (orange-colored adhesions in Figure 6A) showed slow, persistent progression of the cell edge (Figure 6A). On contrary, a cell in which the longest lifetimes are ~10.6 min (yellow-colored adhesions in Figure 6B) showed fast and more fluctuating protrusion behavior (Figure 6B). Accordingly, we sampled the lifetimes of the top 1 percentile of long-living adhesions per movie and obtained the corresponding protrusion velocities and persistent times of the closest edge segments. Plotting edge velocity and persistence as a function of adhesion lifetime showed that cell protrusions with longer mean adhesion lifetimes were associated with faster protrusion velocity but shortened protrusion persistence (Figure 6C and D).

**Figure 6.**
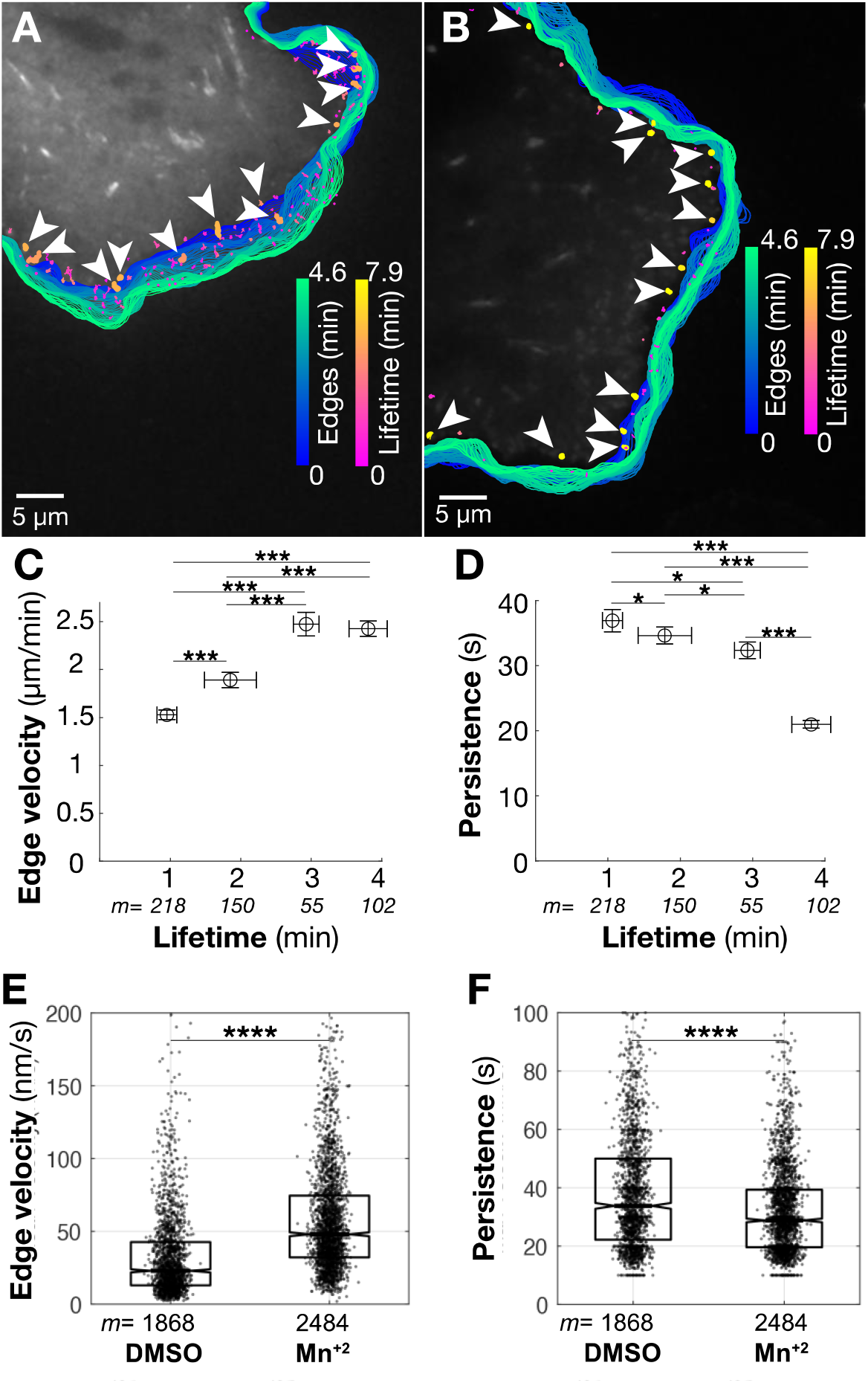
Longer adhesion lifetime is associated with larger edge velocity and shorter persistence. (**A**, **B**) Tracked adhesions and segmented cell edges of COS7 cells with shorter lifetime (A) and longer lifetime (B). Adhesions and cell edges are color-coded for lifetime and frame with Δt = 3 s, respectively. White arrowheads depict longer-living adhesions per each cell movie. (**C**) and (**D**) Error bar plots of velocity (C) and persistent time (D) of edge protrusion in cells with different overall lifetime. Error bar: standard error of mean. * p<0.05, *** p < 10^−15^. *p* value from Man-Whitney’s U test. (**E, F)** Distribution of protrusion velocity (E) and persistence (F), from *m* significant protrusion events in *n* = 7 cells treated with DMSO and *n* = 8 cells with Mn^+2^. Boxes span the 25^th^ to 75^th^ percentile. The central horizontal line is median and notches are 95% CI. *P* value from Kolmogorov–Smirnov test.

We also treated COS7 cells with Mn^+2^, which increases adhesion lifetime and density [44–46, 48]. Mn^+2^ stabilizes nascent adhesions by promoting integrins’ structural shift to high-affinity conformations for binding to extracellular matrix [10, 53, 54]. The cells transiently expressed Emerald-Lifeact to label the cell edge. We imaged the cells’ steady-state protrusion-retraction cycles and quantified protrusion velocity and persistence with morphodynamics software. We found that integrin activation with Mn^+2^ increased mean protrusion velocity but decreased persistence when compared to untreated cells (Figure 6E and F). Together, these results support our model that longer adhesion lifetimes are associated with faster protrusion velocity but reduced protrusion persistence.

## Discussion

Using our novel particle-based BR-MC model, we discovered that actin polymerization is the main driver of lamellipodium velocity and that the force-dependent clutch mechanism of nascent adhesions differentially controls lamellipodium velocity and persistence. Experiments in migrating epithelial cells substantiated that nascent adhesion lifetime promotes protrusion velocity and limits persistence. Directional migration requires persistent edge motion [1, 2], which is optimal at intermediate extracellular matrix density and nascent adhesion concentration [55]. Our findings suggests that in addition to extracellular matrix density, the strength of the adhesion-actin interaction controls protrusion persistence.

Our study clarifies the contributions of lamellipodium actin polymerization and nascent adhesions to overall edge motion. Previous models have indicated both that actin polymerization is sufficient to drive edge protrusion and that adhesion promotes protrusion [28, 31, 56–59]. In the recent model by Garner et al., filament polymerization alone generated stable protrusion [60], which resembles the lamellipodia of fish keratocytes that glide with a static cell shape [61]. Yet, the density of adhesion activation has been shown to promote and be required for protrusion velocity and persistence in protrusion-retraction cycles [28, 59]. We showed that increasing actin polymerization most significantly enhances protrusion velocity and that increasing the nascent adhesion lifetime further supports edge velocity through the molecular clutch mechanism. However, we found that increasing nascent adhesion lifetime reduced the edge motional persistence when membrane tension was moderate, as in the initiation phase of membrane motion. In live cells cycling through the phases of protrusion initiation, reinforcement, and retraction, nascent adhesion lifetime associated with and promoted lamellipodium protrusion velocity but limited persistence. This suggests that the nascent-adhesion mediated regulation in the beginning of edge protrusion dictates the overall protrusion activity.

We propose a mechanism for nascent adhesion lifetime’s differential control of edge velocity and persistence: the increase in edge velocity emerges from the traction that the adhesions exert on the substrate to convert actin retrograde flow into edge motion, while the decrease in persistence arises from the force-velocity relation of actin filaments polymerizing against the plasma membrane. In the initial phase of lamellipodium protrusion, increasing adhesion lifetime tethers the actin filament tips in proximity of the edge, which supports actin filament pushing against the membrane until membrane tension stalls polymerization. The longer the time of filament anchoring, the faster this effect is reached, resulting in decreased protrusion persistence with increasing τ_*max*_. However, when membrane tension has increased, such as at the end of the reinforcement phase, the increase in membrane tension that results from polymerization is too nominal to affect the duration of actin filament polymerization. This model, developed from our computational simulations, is supported by our experimental studies and a complementary study in HT-1080 fibrosarcoma cells migrating slowly on aligned fibers of extracellular matrix [62]. Our tracking of COS7 cell adhesion lifetime and edge motion found that a few long-living adhesions govern edge protrusion. In the HT-1080 study, adhesion lifetime was experimentally controlled by fiber orientation such that adhesions aligned with the matrix had longer lifetimes and more persistent protrusions. When mature adhesions were removed through inhibition of myosin II, the remaining nascent adhesions did not control persistence. Rather, extracellular matrix fibers biased protrusion persistence along the fibers through contact guidance [62].

Our BR-MC model revealed this critical and differential role of nascent adhesions in supporting membrane motion because it incorporated the Brownian ratchet and molecular clutch mechanisms without imposing feedback relations between the actin, adhesion, and membrane dynamics. Prior models of lamellipodial protrusion at the sub micrometer length scale probed the role of actin polymerization using the Brownian ratchet force-velocity relationship of actin filament elongation pushing the membrane [60, 63, 64], or probed actin and adhesion interactions by representing the actin as a gel with the molecular clutch adhesion lifetime-traction force relationship [27, 28, 30]. However, none of the previous models [28, 56, 57, 65–67], to the best of our knowledge, simultaneously incorporated the Brownian ratchet and molecular clutch mechanisms without predetermined relations between membrane motion and actin filament concentration or actomyosin contractility. In our model, lamellipodium dynamics emerge spontaneously from interactions and force balance between actin filaments, membrane, and adhesions. Because we focused on the roles of nascent adhesion dynamics and myosin-independent actin flow, the BR-MC model does not incorporate effects from load adaptations within the actin network or actomyosin contractility. Thus, we cannot exclude that membrane tension feedback to actin network geometry and density [68–71] or traction force feedback to actomyosin contractility and adhesion stabilization [29, 72] influences nascent adhesion regulation of protrusion. Nevertheless, our experiments in cells detected the model’s relationship between adhesion lifetime on protrusion velocity and persistence. This suggests that nascent adhesions also drive lamellipodium velocity and limit persistence in the context of other complex interactions.

In summary, we establish an unexpected role for nascent adhesion lifetime in the differential regulation of protrusion velocity and persistence. We previously showed that integrin subunits have distinct flexibilities and conformations that affect the affinity and avidity of interaction with the matrix [73]. The resulting strength of the integrin-actin molecular clutch interaction controls adhesion lifetime, which controls lamellipodium actin assembly. Thus, while actin polymerization is needed for nascent adhesion formation in edge protrusion [29, 31], our findings indicate that integrin adhesions also control actin assembly. Future studies are needed to determine which integrin-extracellular matrix interactions promote intermediate adhesion lifetime for directed migration in heterogenous environments and how biochemical and mechanical signaling affects adhesion lifetime to alter cell migration.

## Materials and Methods

### Model of lamellipodium protrusions based on Brownian Ratchet and Molecular Clutch (BR-MC)

We developed a 2D model of lamellipodium protrusion based on combining the Brownian ratchet mechanism for actin filament polymerization against a cell edge [11, 22, 35, 37] with the molecular clutch mechanism for adhesions [38, 39]. The model domain is 2 μm x 0.5 μm rectangular domain, with a moving boundary at the top that mimics the flexible cell edge membrane (Figure 1A). Explicit elements in the domain are actin filaments modeled as rigid rods of interconnected units, a flexible membrane of 101 elastically interconnected units, and adhesions modeled as single point particles that create dynamic anchor points for the filaments. Within the domain, actin filaments fluctuate in Brownian motion, polymerize at their barbed ends, depolymerize at their pointed ends, create branches against the membrane and become capped. Actin filaments polymerize against the edge membrane at assembly rate *k_pol_* and branching rate *k*_*branch*_, which generates pushing force that induces outward membrane motion (Figure 1A). Integrins undergo cycles of activation and deactivation, which correspond to their addition to and removal from the simulation domain. These mechanisms are governed by kinetic rates (Table 1). Filaments and integrins interact by establishing potential energies, and filaments push the edge membrane by exerting force on it. The membrane motion increases membrane tension, ***F***_*M*_, which results in retrograde flow of the actin filaments. Through the retrograde flow, the ***F***_*M*_ is transmitted to the adhesions in the form of traction force. In response to the traction force, adhesions dynamically unbind at a force dependent rate, *k*_*off*_. Displacements of actin filaments and membrane are calculated over the course of the simulation as a function of the forces acting on them, using the overdamped Langevin Equation as:

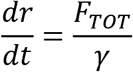

where *F*_*TOT*_ is the sum of deterministic and stochastic forces, *r* is a position vector for the elements in the system, *γ* is the environment drag coefficient (assuming cytoplasm viscosity), and *dt* is the simulation time step. A complete description of the model, including the implementation scheme and relative interactions between actin, adhesions and membrane, is below.

#### Model initialization

During the initialization of the model, a membrane is placed on the top boundary and the actin filaments are randomly distributed within the domain. Each actin filament is initialized with a length of *n* units, with each unit representing 5 monomers or 15 nm. The *n* number of units is randomly selected from a normal distribution with a mean of 15, corresponding to an average filament length of 225 nm, a standard deviation of 2, and a minimum *n* of 1. The concentration of actin filaments is maintained at a steady state of 9000 filamentary units in the simulation domain. This creates a buildup of actin filaments during the first second of simulations, with the occasional addition of filaments thereafter in order to maintain a steady concentration of filaments.

A finite number of integrins are allowed to activate and deactivate according to their kinetic rates, until they reach a steady state concentration. Allowed adhesions = 1000 when simulations are run using a force-independent *k*_*off*_. Allowed adhesions = 2000 with force-dependent *k*_*off*_. The effective concentration of integrins emerges in the model from the relative magnitudes of *k*_*off*_ and *k*_*off*_.

#### Model iterations

The model is developed using MATLAB R2020b, using the explicit Euler implementation scheme for performing each model iteration, which consists of kinetics, force balance, and position updates. The kinetic events occur following sequential steps: polymerization, depolymerization, capping and branching of the actin filaments, and adhesion activation and deactivation. Then, interactions between actin filaments and membrane and between filaments and adhesions are evaluated and the corresponding forces are calculated. Last, based on the force balance between interacting elements, the relative positions between actin filaments and membrane, and between actin filaments and adhesions are calculated, and the boundary conditions applied.

#### Actin filament representation and dynamics

In the model, actin filaments are represented as rigid and polar rods of interconnected units (Figure 1A). Their polymerization occurs as addition of units at the filament barbed ends, with rate *k_polym_*. Depolymerization occurs as removal of units from the pointed ends, at a rate *k_depolym_*. Actin filaments within a branch window of 15 nm from the membrane can also form branches at a rate *k*_*branch*_. Branching occurs as elongation of new filaments from existing (mother) filaments in the direction towards the membrane and at an angle *θ* relative to the mother filament. *θ* is randomly selected from a normal distribution with a mean of 70 deg and standard deviation of 10 deg. Filaments capping occurs at a rate *k_cap_*, which stops filaments polymerization and branching.

#### Actin filament polymerization follows the Brownian ratchet mechanism

In order to account for the effects of membrane tension on actin filaments polymerization, filaments presenting barbed ends within 15 nm from the membrane slow their polymerization rate depending on the force on them. For these filaments, the probability of polymerization is calculated as *P*_*polym*_ = *C*_*p*_ *k*_*polym*_ *dt,* where the polymerization coefficient, *C*_*p*_, varies between 0 and 1:

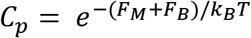

where *F*_*M*_ is membrane tension, *F*_*B*_ is a confining boundary which ensures a reflective boundary, and *k*_*B*_*T* = 4.11 pN nm.

#### Actin filament connection to adhesions

When a filamentary unit is within a distance *r*_*thresh*_ from integrin, a harmonic interaction potential is established between the filament and integrin, with force: *F*_*A*_ = *r*_*0,A*_ *k*_*A*_, where *k*_*A*_ is the integrin-filament spring constant, and *r*_*0,A*_ is the distance from the equilibrium, resulting from actin retrograde flow. When an integrin switches its state from active to inactive, the connection with the actin filament is lost and integrin disappears.

#### Filament motion follows Langevin dynamics

The total force on each actin filaments is computed as: *F*_*TOT*_ = *F*_*T*_ + *F*_*A*_ + *F*_*M*_ + *F*_*B*_, where *F*_*T*_ is a Brownian, stochastic force following the fluctuation dissipation theorem, *F*_*A*_ is the force from one or more bound integrins, *F*_*M*_ is the force exerted by the membrane, and *F*_*B*_ is the boundary force which acts as a repulsive potential preventing the filaments from crossing. When one filament presents one or more branches and/or branches on branches, the interconnected filaments are treated as a rigid structure.

Filaments positions are calculated over time following Langevin equation in the limit of high friction: 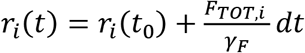 *for i* = 1,2,3, … *N*, where *N* is the total number of filaments and filament structures, *γ*_*F*_ is the frictional coefficient and *dt* is the simulation time step.

#### Integrin representation and dynamics

Integrins are represented as single point particles existing in two functional states: active or inactive, as they undergo cycles of activation and deactivation at rates *k*_*on*_ and *k*_*off*_, respectively. When active, integrins are placed in random positions in the simulation domain and provide anchor points for filaments motion. When inactive, they are removed from the domain. While integrin activation is governed by *k*_*on*_, their de-activation occurs through one of two mechanisms: force-independent or force-dependent unbinding. In the first case, *k*_*off*_ has a constant value (Table 1). In the second case, *k*_*off*_ depends biphasically on the tension between actin filament and integrin, *F*_*A*_. According to the catch-bond model for integrin unbinding, the unbinding rate is a sum of two exponentials with opposite signs (Figure 4A):

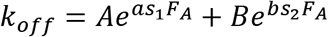

Three lifetimes versus force relations were used in the model (lifetime τ = *1/k_off_*) and they are here referenced according to their maximum τ values (τ_max_):

| | $\tau_{max} = 3$ | $\tau_{max} = 7.5$ | $\tau_{max} = 12$ |
| --- | --- | --- | --- |
| A | 2.0 | 0.5 | 0.25 |
| B | 0.5E-5 | 0.5E-5 | 0.5E-5 |
| a | -9.727E-3 | -7.286E-3 | -6.12E-3 |
| b | 0.3 | 0.3 | 0.3 |
| $s_1$ | 6.8 | 6.8 | 6.8 |
| $s_2$ | 1.0405 | 0.9228 | 0.8606 |

#### Membrane representation and dynamics

The edge membrane is represented with consecutive rigid rods, each 18 nm long, interconnected by harmonic interaction potentials of stiffness *k*_*m*_ and equilibrium separation *r_0,M_*. Each membrane segment experiences two forces: the filaments pushing against it, and the pulling from neighboring membrane segments.

Over the course of the simulations, positions of membrane segments are updated following the Langevin equation, as: 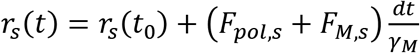 *for s = 1,2,3,…101,* where *F*_*pol*_ is the force from filaments, and *F*_*M*_ is the elastic force contribution from the connection with neighboring membrane segments. *γ*_*F*_ is the drag coefficient from the fluid in front of the membrane (outside the cell).

#### Determination of traction force

Mean traction force is the average of all adhesion-filament connections over all time points and positions. All force values greater the 500 pN resulted from motion too fast for the 1 ms timestep and were excluded (*F_A_* < 500 pN).

#### Extraction of protrusion persistence

Velocity data for each membrane segment was decomposed into its Intrinsic Mode Functions (IMF) using Empirical Mode Decomposition. The first 7 IMFs were subtracted so that thermal noise oscillations in membrane velocity were on the scale of real cell protrusion oscillations, in which change in velocity is at least 5 nm/s (Figure 5 - figure supplement 1).

### Key resources table

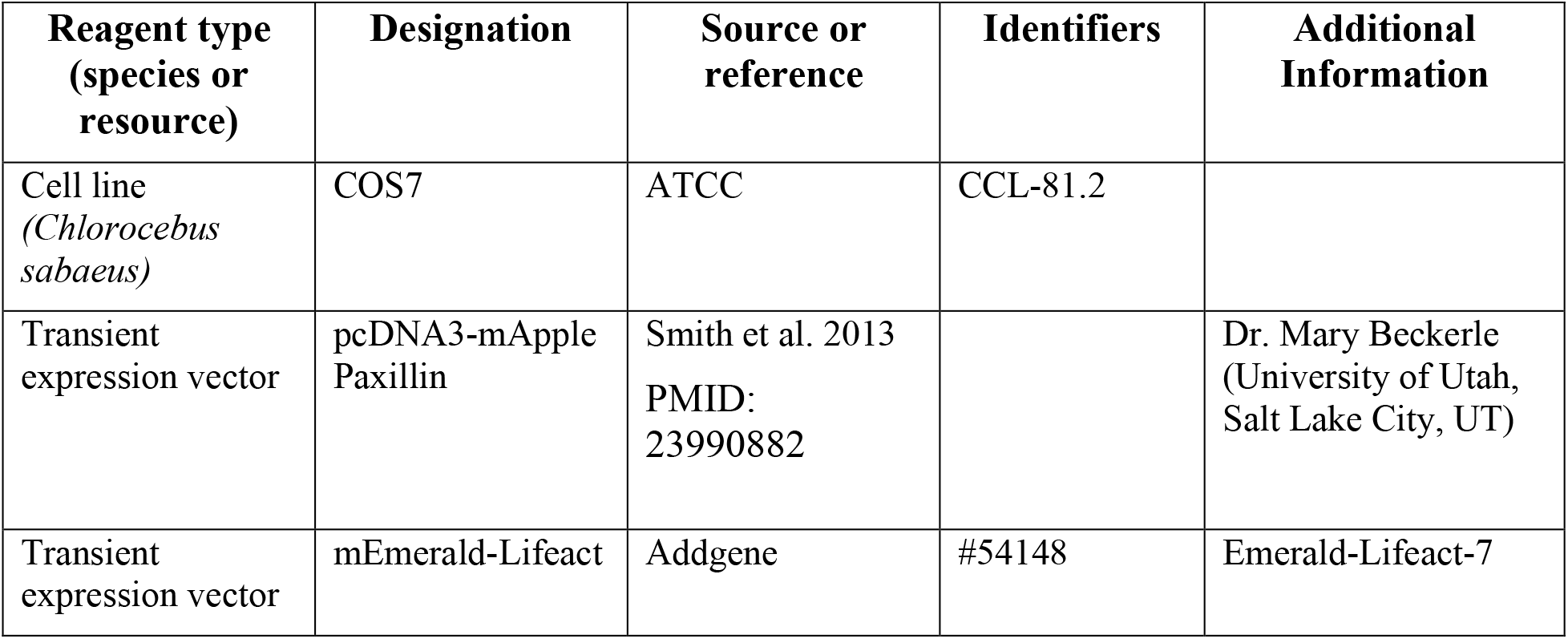

| Reagent type<br>(species or<br>resource) | Designation | Source or<br>reference | Identifiers | Additional<br>Information |
| --- | --- | --- | --- | --- |
| Cell line<br>(*Chlorocebus*<br>*sabaeus*) | COS7 | ATCC | CCL-81.2 |  |
| Transient<br>expression vector | pcDNA3-mApple<br>Paxillin | Smith et al. 2013<br>PMID:<br>23990882 |  | Dr. Mary Beckerle<br>(University of Utah,<br>Salt Lake City, UT) |
| Transient<br>expression vector | mEmerald-Lifeact | Addgene | #54148 | Emerald-Lifeact-7 |

### Cell culture

COS7 cells were obtained from ATCC, cultured in Dulbecco’s Modified Eagle Medium (DMEM) with 4.5 g/L D-glucose, L-glutamine, and sodium pyruvate (Gibco 11965092) containing 5% Fetal Bovine Serums (FBS, Avantor Seradigm 97068-085), and tested for micoplasma every 3-6 months. Micoplasma-negative COS7 cells were plated on acid-treated 1.5 coverslips within 35 mm glass-bottom dishes (MatTek P35G-1.5-14-C), and transfected the following day with pcDNA3.1/Paxillin-mApple or mEmerald-Lifeact at 20% confluency using TransIT-LT1 (Mirus MIR 2304) following the manufacturer’s instructions. Two days post-transfection, medium was replaced with FluoroBrite DMEM (Gibco #) supplemented with 10% FBS and 20 mM HEPES.

### Live cell TIRF imaging of adhesions

For adhesion imaging, cells were imaged on an automated Nikon Ti inverted microscope with motorized total internal reflection fluorescence (TIRF), Perfect Focus 3 to maintain laser-based identification of the bottom of the substrate during acquisition, a CFI Apo TIRF 100x oil Apo 1.49 NA objective, 561 solid-state laser (Vortran), ET620/60m emission filter (Chroma), and Photometrics Prime 95B camera configured at a 100 MHz readout speed to decrease readout noise with Metamorph. Images were taken every 3 s for 5 min, with sequential images at every time point with the TIRF angle set to optimal TIRF and with the TIRF angle set as vertical for effective widefield imaging. The acquired images had an effective pixel size of 45 nm. Imaging was performed at 37°C, 5% carbon dioxide, and 70% humidity. Laser powers were decreased as much as possible and the exposure time set at 200-400 ms to avoid phototoxicity.

### Adhesion segmentation, detection, and tracking

Nascent adhesions were detected and segmented using point source detection as previously described in [51, 74]. Briefly, fluorescence images were filtered using the Laplacian of Gaussian filter and then local maxima were detected. Each local maximum was then fitted with an isotropic Gaussian function (standard deviation: 2.1 pixels, i.e. ~180 nm) and outliers were removed using a goodness of fit test (p=0.05). The point sources detected for nascent adhesions were tracked over the entire frames of the time-lapse images using uTrack [75]. Lifetimes of adhesions were calculated from lifetimes of individual tracked trajectories. A GUI-based MATLAB software for the edge analysis is from Danuser lab [52]. Due to the noise in edge motion from the 3 s framerate, IMFs were removed in the edge motion data.

### Live cell confocal imaging of cell edge and analysis

For cell edge imaging, cells expressing mEmerald-Lifeact were imaged on a Nikon Ti inverted microscope with a CFI Apo TIRF 60x oil, 1.45 NA objective employing Perfect Focus, a Yokagawa CSU-10 spinning disk confocal with Spectral Applied Research Borealis modification, a 488 solid-state laser, Chroma ET525/50m filter, and Photometrics Myo CCD camera with Metamorph. Images were taken every 10 s for 5 min, with 400-700 ms exposures. For experiments with prolonged adhesion lifetime, cells were treated with 1 mM MnCl_2_ 2 h prior to imaging. Time-lapse images were analyzed for cell edge protrusion dynamics in MATLAB software as described previously [76]. A GUI-based MATLAB software for the edge analysis is from Danuser lab [52]. A two-sample nonparametric Kolmogorov–Smirnov test at 5% significance tested for population distribution equality. IMFs were not removed from this edge motion data.

### Statistics

Models were run until additional iterations no longer changed the result output. The non-parametric Mann-Whitney U test was used to test for difference in the means for all modeling data and adhesion-edge analyses. Experiment sample size was chosen based on a minimum of three independent biological replicates and hundreds to thousands of adhesions and protrusion events analyzed, respectively, within each replicate. The Kolmogorov-Smirnov test was used to test for difference in the distribution of edge motion upon Mn+2 treatment. Unless otherwise mentioned in each figure caption, * *p*<0.05, ** *p*<0.01, *** *p*<0.001, **** *p*<0.0001.

### Software availability

MATLAB software for the computational model is shared via GitHub at https://github.com/KRCSLC/ProtrusionModel.

A GUI-based MATLAB software for the adhesion analysis is shared via GitHub at https://github.com/HanLab-BME-MTU/focalAdhesionPackage.git; Han,2021; copy archived at swh:1:rev:6aeb3593a5fd3ace9b0663d1bf0334decfb99835.

## Acknowledgements

Thanks to Drs. Mark Smith and Mary Beckerle for the gift of pcDNA3-mPaxillin-mApple. Thanks to Dr. Drew Elliott for acquisition of the timelapse images of Paxillin-mApple in COS7 cells. Thanks to the University of Utah Cell Imaging Core and the University of Utah Center for High Performance Computing. The work was supported by funding from the National Science Foundation NSF-BMMB-2044394 to T.B.D., the Huntsman Cancer Institute and Scientific Computing and Imaging Institute CORI to T.B.D., American Cancer Society RSG CSM130435 to M.C.M. and National Institutes of Health 1R15GM135806 to S.J.H.

## Competing Interests

The authors declare no competing interests.

## Notes

### Competing Interest Statement

The authors have declared no competing interest.

